# Developmental regulation of Canonical and small ORF translation from mRNAs

**DOI:** 10.1101/727339

**Authors:** Pedro Patraquim, Muhammad Ali S. Mumtaz, Jose I. Pueyo, Julie L. Aspden, J.P. Couso

## Abstract

Ribosomal profiling has revealed the translation of thousands of sequences outside of annotated protein-coding genes, including small Open Reading Frames of less than 100 codons, and the translational regulation of many genes. Here we have improved Poly-Ribo-Seq and applied it to *Drosophila melanogaster* embryos to extend the catalogue of *in-vivo* translated small ORFs, and to reveal the translational regulation of both small and canonical ORFs from mRNAs across embryogenesis. We obtain highly correlated samples across five embryonic stages, with close to 500 million putative ribosomal footprints mapped to mRNAs, and compared them to existing Ribo-Seq and proteomic data. Our analysis reveals, for the first time in *Drosophila*, footprints mapping to codons in a phased pattern, the hallmark of productive translation, and we propose a simple binomial probability metric to ascertain translation probability. However, our results also reveal reproducible ribosomal binding apparently not resulting in productive translation. This non-productive ribosomal binding seems to be especially prevalent amongst upstream short ORFs located in the 5’ mRNA Leaders, and amongst canonical ORFs during the activation of the zygotic translatome at the maternal to zygotic transition. We suggest that this non-productive ribosomal binding might be due to cis-regulatory ribosomal binding, and to defective ribosomal scanning of ORFs outside periods of productive translation. Finally, we show that the main function of upstream short ORFs is to buffer the translation of canonical ORFs, and that in general small ORFs in mRNAs display Poly-Ribo-Seq and bioinformatics markers compatible with an evolutionary transitory state towards full coding function.

## BACKGROUND

Ribosomal profiling, the genome-wide sequencing of ribosome-protected RNA fragments (Ribo-Seq) has been increasing our understanding of a crucial event in all living genomes: protein translation [1, 2]. Ribo-Seq results show that there exists a noticeable level of non-canonical translation in eukaryotes. This can arise from non-AUG codons, from polycistronic transcripts, and from unannotated small Open Reading Frames of less than 100 codons (smORFs or sORFs) [3–5].

At the same time, the comparison of RNA-Seq and Ribo-Seq for the same biological samples allows the transcriptome-wide study of an important but often forgotten regulatory process in genome function: the regulation of translation. Typically, expression of an mRNA transcript is equated with the automatic translation of any encoded canonical ORF (of more than 100 codons). However, translation is not automatic, but is regulated, often with great relevance for organismal function, both under normal and altered conditions [6–10]. Translation regulation is also a key aspect of development in all animals [11] and has been studied extensively in *Drosophila* [12–14]. *Drosophila* embryogenesis is a highly-coordinated and complex process that is completed in a time span of just 24 hours [15]. During the first two hours after egg laying (AEL) there is absence of transcription from the zygotic genome and the key developmental processes, such as establishment of the primary Antero-Posterior and Dorso-Ventral axes, are controlled purely through the translational regulation of maternal mRNA previously laid in the egg [16, 17]. After this initial period, the embryo undergoes a ‘maternal to zygotic transitioń, whereby the transcription and translation of the zygotic genome takes over the maternal products, a process also found in nematodes, echinoderms, and vertebrates [13, 18–20]. Nonetheless, the impact of translational regulation at the genome-wide scale on the whole of *Drosophila* embryogenesis has not yet been revealed.

Ribo-Seq results regarding non-canonical and regulated translation have been the subject of debate. While it has become accepted that both processes may occur more extensively than previously thought, there is no consensus on the actual fraction of smORFs and non-canonical ORFs whose translation is shown by Ribo-Seq [21–25]. The Ribo-Seq debate centres on the asymmetry between these numbers and other translational evidence, and on the interpretation of the Ribo-Seq results themselves. The most widely used counterpart of Ribo-Seq is proteomics, but the numbers of protein and peptides detected by proteomics consistently fall short of those detected by Ribo-Seq, especially regarding non-canonical translation. For example, the most thorough proteomics study to date covering the whole *Drosophila melanogaster* life-cycle has detected less than 40% of all unique canonical proteins [26]. This number is further reduced to 30% of annotated smORF polypeptides, while we have previously reported that 80% of canonical and small ORFs show clear Ribo-Seq evidence of translation in a single embryonic cell line [23]. However, Ribo-Seq detects ribosomal binding, not actual peptide production. There is not a universally agreed Ribo-Seq metric unequivocally identifying productive, biologically-relevant translation, as opposed to other processes such as low-level background translation, ribosomal scanning and nonsense-mediated-decay surveillance, or stochastic ribosomal binding. Bioinformatically, it is accepted that ribosomal binding above a certain level, and especially, binding showing tri-nucleotide periodicity in phase with codon triplets (phasing or framing), indicate translation of an ORF [1, 2]. A biochemical approach is to introduce modifications to the ribosomal-RNA purification, to ensure that only ribosomes engaged in productive translation are selected. For example Ribo-Seq of polysomes (RNAs bound by several ribosomes), since the sequential translation of polyadenylated, capped and circularised mRNAs by several ribosomes is a supramolecular feature of productive translation, and excludes single ribosomes (which could be involved in low-level translation but also in other activities) [23, 27]. We have called this latter approach Poly-Ribo-Seq [23, 27].

Here we present an *in vivo* Poly-Ribo-Seq study covering a time-course of *Drosophila melanogaster* embryogenesis. We have both improved our experimental Poly-Ribo-Seq and the subsequent data analysis pipeline, to obtain unprecedented levels of Ribo-Seq efficiency (reads mapped to ORFs) and quality, including codon framing as the hallmark of productive, biologically meaningful translation. Thus, we can ascertain translation and its regulation *in vivo* and across development for both canonical and non-canonical ORFs. We detect the translation of thousands of non-annotated ORFs, and identify hundreds of mRNAs whose translation is highly regulated during embryogenesis. However, our results also reveal reproducible ribosomal binding not resulting in productive translation. This non-productive ribosomal binding seems to be especially prevalent amongst upstream short ORFs located in the 5’ mRNA Leaders, and amongst canonical ORFs during the activation of the zygotic translatome at the maternal to zygotic transition. We suggest that this type of ribosomal binding might be due to either cis-regulatory ribosomal activity, or to defective ribosomal scanning of ORFs outside periods of productive translation.

## RESULTS

### 1) The method and overall data

Since Poly-Ribo-Seq requires even larger amounts of starting material than Ribo-Seq due to polysome fractionation, the *Drosophila S2* cell line was an excellent tool for the development of the technique. However, the S2 cell line is derived from just one type of tissue (macrophage-like) from late stage *Drosophila melanogaster* embryos [28] and may not be the ideal system to obtain a comprehensive picture of *Drosophila* translation. Only 60% of both canonical genes and uORFs, and only about a third of annotated smORFs (hereafter referred as short CDSs, or sCDSs [29]) appear transcribed in this cell line [23]. Therefore, we applied Poly-Ribo-Seq to *D. melanogaster* embryos in order to extend the catalogue of *in-vivo* translated smORFs and study translation across three distinct developmental time windows of *Drosophila* embryogenesis. The original protocol from Aspden [23] was adapted and improved in terms of biochemical rRNA depletion and bioinformatics mapping of FPs to ORFs (see methods), and we obtained higher yield, purity and accuracy (Fig.1 and Sup. Table 1). This way we detected rare but reproducible translation events in non-annotated genes, and obtained a comprehensive picture of translation across development.

**Figure 1.**
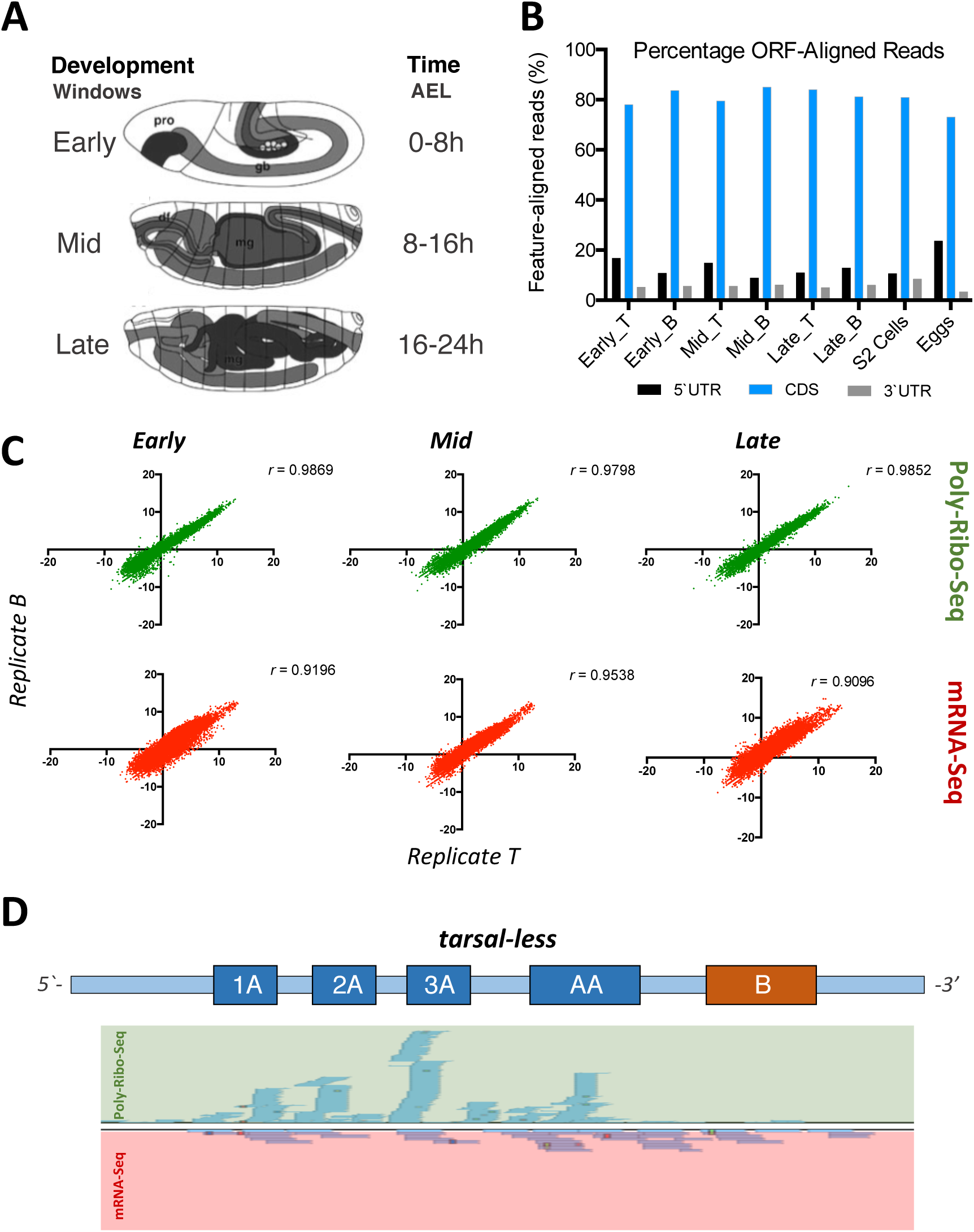
Experimental design and quality-control of Poly-Ribo-Seq data. **A** *Drosophila melanogaster* embryos were collected in three contiguous 8 hour time-windows spanning the whole of embryogenesis. Two biological replicates were collected for each window, for both Poly-Ribo-Seq and RNA-Seq. pro: procephalon; gb: germ band; df: germ band; mg: presumptive midgut **B** Percentage of 5’UTR, 3’UTR and CDS-aligned reads for Poly-Ribo-Seq, across the annotated transcriptome of *Drosophila melanogaster*. **C** Reproducibility of RPKM values for canonical ORFs by Spearman correlation (*r*) across replicates T (x-axis) and B (y-axis), per stage, for both Poly-Ribo-Seq (top panels) and RNA-Seq (bottom panels). **D** Density of genome-aligned reads for all experiments, both Poly-Ribo-Seq and RNA-Seq across the polycistronic *tarsal-less* (*tal*) transcript.

In our experimental design, we divided the *Drosophila* embryogenesis in three temporal 8 hour windows: Early (0-8 hours after egg laying, or 0-8h. AEL), Mid (8-16h. AEL) and Late embryogenesis (16-24h. AEL) (Fig. 1A). The key developmental processes occurring during these periods of embryogenesis are described elsewhere [15]. Briefly, early embryogenesis (0-8h. AEL) is characterised by the maternal to zygotic transition, followed by gastrulation, germ band formation and ectodermal and mesodermal determination and pattern formation, including the determination of the CNS. The 8-16h. AEL hours of development (mid-embryogenesis) covers the process of tissue formation and organogenesis (such as presumptive gonad and gut assembly) including multi-organ processes such as formation of segments and the feeding cavity. The differentiation of neurons and muscles also occur during this time. Late embryogenesis (16-24h. AEL) is characterised by final differentiation and the start of physiological processes such as epidermal and cuticle differentiation [30] (with opening of tracheae to respiration) and digestive system differentiation (with yolk digestion and uric acid) [31, 32], plus the connection and fine-tuning of neural, sensory and muscle structures, eventually resulting in coordinated embryo movements and larval hatching [33, 34].

We collected two biological replicas (called T and B) from each embryonic stage, extracting simultaneously total RNA and Ribosomal Footprints. Sequencing and analysis of these samples was used to determine transcription and translation levels, which in turn were used to reveal the extent and quality of translational regulation during *Drosophila* embryogenesis at a genomic scale. We studied several classes of ORFs as defined in Couso and Patraquim [29], altogether numbering 40,852 ORFs (see methods): 21,118 canonical ORFs of more than 100 codons in polyadenylated mRNAs; 862 ORFs from annotated coding sequences of 100 codons or less in an otherwise canonical mRNA (short CDSs); and 18,872 upstream ORFs (uORFs) in the 5’ leaders of canonical mRNAs. We studied only AUG-START ORFs, and also discarded overlapping uORFs, in order to ensure the accuracy of our translation assessments. Similarly, the translation of ORFs in putative long-non-coding RNAs (lncORFs) will be described in a forthcoming study due to its special characteristics.

As the Ribo-Seq protocol typically results in a composite library with a majority of ribosomal rRNA reads, and a minority of ribosomal footprints (FPs), we sequenced an exhaustive amount, totalling 1.2 billion Ribo-Seq reads, yielding some 345 million ribosomal footprints (FPs) mapped to the genome (Sup. Table 1) with ∼80% FPs aligned to annotated CDS, ∼16% to 5’ Leaders and 6% to 3’UTRs (Figs.1B and 2A), providing a 300x coverage or our 40,852-strong ORF-ome. The data shows high reproducibility, with high Spearman correlation(*r*=0.9) of RPKMs (number of sequenced reads per ORF length in kilobase, per million reads) for each ORF amongst the two replicas (Fig. 1C) for both RNA-Seq (RPKM^RNA^) and Ribo-Seq (RPKM^FP^) samples. Only reads of 26 to 36 nucleotides (nt) were counted for RPKM^FP^, as this length distribution corresponded to 97% of the reads obtained in all cases, and is within the size range previously described for Ribosomal FPs [1] (Sup. Fig 1A). At first glance, our data reveals translation of non-canonical genes, as for the polycistronic smORF gene *tarsal-less* [35] (Fig. 1D and Sup. Table 3A).

**Figure 2.**
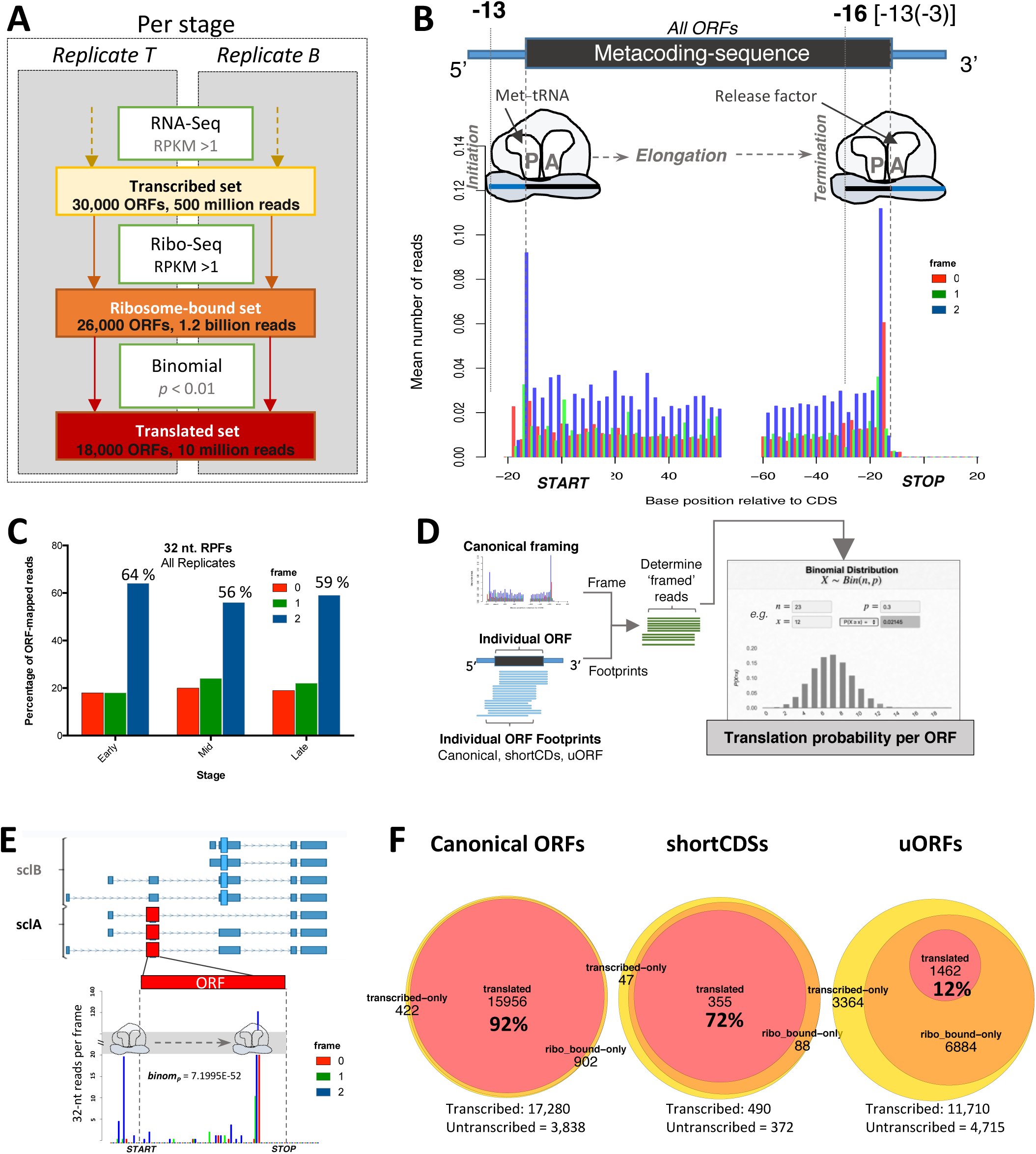
Detection of Translation in annotated and non-annotated ORFs. **A** Sequentially inclusive pipeline used for translation calls. An ORF was considered translated, in each stage, if it showed RPKM > 1 in one of two RNA-Seq replicates, RPKM > 1 in both Poly-Ribo-Seq replicates and a *p*-value < 0.01 in the binomial test for framed-reads. **B** Metagene plot for codon framing of 32-nt. ribosome footprints across all ORFs analysed in this study, showing three-nucleotide periodicity (framing) in the third position of each codon (frame 2, blue). Numbers on top denote the distances between the 5’-end of ribosomal footprints and START/STOP codons. **C** The observed framing for 32-nt. Ribosome footprints is consistent across embryonic stages (combined replicates per stage). **D** Logic of per-ORF analysis of framing probabilities using the binomial test. **E** Density of 32 nt. genome-aligned Poly-Ribo-Seq reads for all experiments, per frame, and corresponding translation probability for the *scl-A* ORF, as measured by the binomial test. **F** Numbers of developmentally-transcribed, ribosome-bound and translated canonical ORFs (annotated ORFs with length > 100 codons), shortCDSs (annotated ORFs ≤ 100 codons) and uORFs.

### 2) Framing: an indicator of productive translation

For our bioinformatic analysis (see Methods and Fig. 2A), we use RPKM^RNA^>1 to indicate transcription of an ORF (in either of our two replicas). To determine translation, we use two combined filters. First, a quantitative filter: a minimal amount of ribosomal binding must be detected, set as a standard RPKM^FP^>1 in both T and B replicas, to indicate signal above error and background. Second, a qualitative filter, the nature of that binding: the ribosomal movement across an ORF during translation generates tri-nucleotide phasing of ribosomal protected footprints, or framing (Fig. 2B), in contrast with the reads derived from RNA-Seq, which accumulate evenly across all nucleotides in mRNA (Sup. Fig 1B).

Previous ribosomal profiling studies of *Drosophila* had not been able to observe strong framing on a genome-wide scale, and thus translation has been proposed on the basis of RPKM and other quantitative metrics, such as coverage [18, 23, 24, 36]. However, our improved Poly-Ribo-Seq method, coupled with an improved bioinformatics analysis of the data (see methods), revealed framing for the first time in the *Drosophila* (Figure 2B-C). Like all Ribo-Seq experiments, our data revealed a distribution of Ribosomal footprint lengths (Sup. Fig 1A). We focused on the most represented read length (32 nucleotides), which also shows the clearest framing in all samples (55-60% reads in-frame; Figure 2C). Different formulae and metrics have been previously used to quantify framing per ORF and provide a cut-off [1, 21], reviewed in [5, 37], which was usually obtained by sampling a small number of bona-fide translated ORFs and extrapolating the results. While these methods can identify translation, they can be complex and difficult to apply universally to all Ribo-Seq datasets, due to differences in sampling and protocols utilised.

Here, we have opted for a simpler approach that is easy to extend to past and future datasets. We treat framing as a simple coin-toss problem, which offers clear statistical probabilities, even with a low number of reads. As each footprint is the result of an independent RNA-ribosome molecular interaction, each resulting sequencing read has in principle a 1/3 probability of aligning to the right frame in an ORF, and 2/3 of aligning to the wrong frame (Fig. 2D). The probability of having *x* successful outcomes (reads aligned to the ORF frame) following *n* independent trials (number of reads mapping to the ORF) follows a classical binomial distribution, which can be consulted using widely available tables [38] and computer programs. We take the observed binomial probability as the probability of being translated for a given ORF and we consider it as indicative of translation when p<0.01 (i.e. the probability of observing such framing of reads in the ORF by chance is less than one in a hundred; see Methods for details and implementation). Here, because of our RPKM^FP^>1 filter, and because only 32 nt. FP reads were used to calculate framing (see methods and Sup.Fig.1) we noted that, out of 44,067 translation calls in all stages studied, only 36 calls appeared translated having less than 20 ribosomal footprints (of 26 to 36nts) and only 2 calls had less than 10 footprints. As example (Fig. 2E and Sup. Table 3; see also Sup. Data for full sets of data) we show the non-canonical smORF gene *sarcolamban* (*scl*) whose translation has been determined experimentally [39]. Its two ORFs (*SclA* and *SclB*) achieve RPKM^FP^ above 1 in both T and B replicas during Mid and Late embryogenesis. Most of its FP reads mapped to frame “2”, but *SclB* only achieved a binomial *p*-value below 0.01 during Late embryogenesis. Thus, our data indicated that *SclA* is ribosomal-bound and translated at Mid- and Late embryogenesis, but *SclB* is only productively translated during Late embryogenesis (when *scl* function in regulating muscle contraction [39] would have its maximum requirement, see above and Fig.1A). Note that framing *p*-values do not indicate the amount of translation (which is indicated by the RPKM^FP^ and its associated metric translational efficiency, TE, see below) but only the likelihood that the observed ribosomal binding leads to productive ORF translation. This binomial framing metric also ensures that the translation detected belongs to a given ORF, not to another putative ORF overlapping it.

Using these criteria, we observed that 98% of transcribed canonical ORFs, 90% of short CDSs and 71% of uORFs were reproducibly bound by ribosomes at any point during development (showed RPKM^FP^>1 in both replicates of the same stage), whereas the percentage of actually translated ORFs (ribosomal-bound ORFs also having framing p-value<0.01 in that stage) was 92% of Canonical, 72% of short CDSs, and 12% of transcribed uORFs (Fig. 2F).

### 3) Consistency of Transcription and Translation measurements across different types of genomic data

We further verified the robustness and reproducibility of our data and analysis by cross-comparisons across related gene-expression genomic data: RNA-Seq, Ribosomal profiling, and quantitative proteomics. For RNA expression, we consolidated the modENCODE embryo RNA-Seq data [40], into our three 8-hour long developmental time windows. For all three developmental windows, the RPKM^RNA^ of RNA-Seq samples was more highly correlated (in the region of *r*=0.9) with a replica of their stage (either ours or modENCODE), than with samples of other stages (Fig. 1C, Fig. 3A, Sup Table 2).

**Figure 3.**
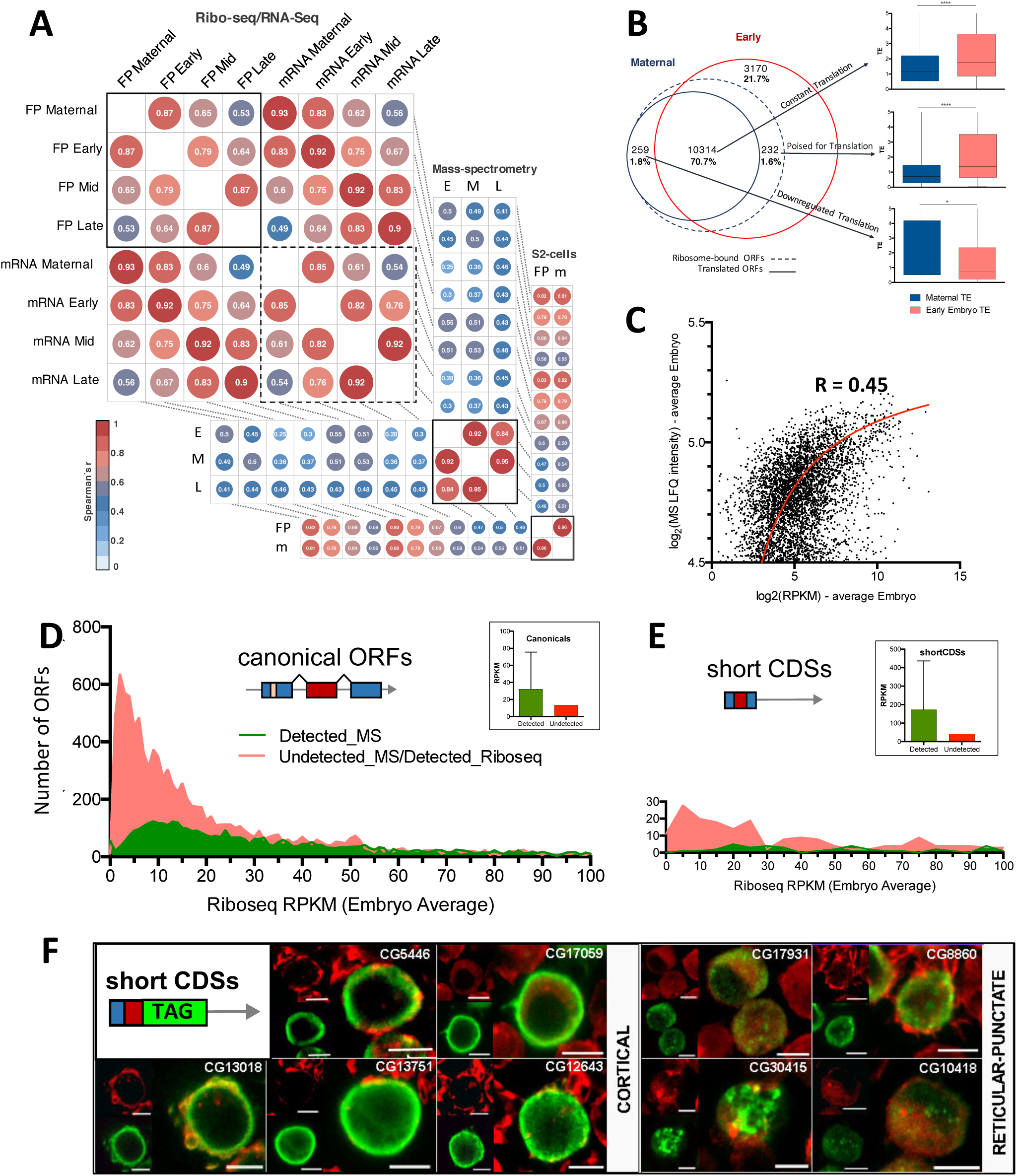
Comparisons across genome-wide datasets. **A** Per-stage Spearmańs correlations across Poly-Ribo-Seq (“FP”), RNA-Seq (“mRNA”) and mass-spectrometry datasets (“E”-Early, “M”-Mid, “L”-Late stages correspondence in *Casas-Vila* et al. 2017 - see Methods). “Maternal” datasets correspond to concatenated Mature Oocyte and Activated Egg datasets from *Kronja et al. 2014*. S2-cells correspond to the *Aspden et al. 2014* dataset plus of novel sequencing. Numbers denote Spearman’s *rho*. **B** Numbers and translation fates of Maternal ribosome-bound canonical ORFs in the maternal-to-zygotic transition. “Constant translation” denotes ORFs detected as translated by our pipeline in both Maternal and Early embryo datasets (see Figure 2A). “Poised for translation” denotes ORFs showing Maternal ribosome-binding which only appear as translated in the Early embryo. “Downregulated translation” denotes ORFs with Ribosome-binding and translation in the Maternal dataset, which do not appear in the Early embryo dataset. **C** Overall correlation between average embryo Poly-Ribo-Seq RPKM and average embryo Mass-Spec imputed LFQ intensities (both datasets are log_2_-transformed). All detected ORFs are included in this analysis. **D** Differential detectability of canonical ORFs between Poly-Ribo-Seq and Mass-spectrometry techniques. **E** Differential detectability of shortCDSs between Poly-Ribo-Seq and Mass-spectrometry. **F** S2-tagging experiments of 9 uncharacterized short CDSs showing translation in either cortical or reticular-punctate patterns (FLAG antibody: green, F-actin stained with phalloidin: red).

For Ribosomal profiling, our RPKM^FP^ data showed an even more marked trend to correlate preferentially first with their own stage replica (r=0.98-0.99), and second with samples from adjacent time periods (Fig. 1C, 3A, Sup Table 2). Our FPs and RNA-Seq from the same stage showed a high correlation of r=0.8-0.9 (Fig. 1C, Sup Table 2). We also compared our FP data with the Kronja et al. 2014 Ribo-Seq data from unfertilized *Drosophila* eggs, that was obtained after RNaseI digestion akin to our own protocol. As expected, the RPKM^FP^ from unfertilized eggs (FP Maternal) showed the highest correlation with our RPKM^FP^ samples from early embryogenesis (0-8h AEL) (Spearmańs *r*≈0.8) (Fig. 3A and Sup. Table 2). Our bioinformatic pipeline detected framing in Kronjás ORF-mapping reads of 29nt (see methods). 10,314 canonical ORFs appeared translated both in unfertilized eggs and Early embryos, albeit showing increased translation efficiency in the latter (Fig. 3B). However, differences between the maternal and zygotic translatomes were also identified, since the Kronja [18] data only assessed maternal RNA translation while our 0-8h AEL sample detects both maternal and early zygotic translation (see “Regulation of translation” below).

We also compared our embryo FP data with *Drosophila* embryo-derived S2 cell cultures [28], which in principle should provide synchronic data less subjected to experimental and developmental noise. We have re-analysed the data from Aspden et al. 2014, adding a new Poly-Ribo-Seq sample obtained here. S2 translation in general is highly correlated with Early embryos (Fig. 3A), and the proportions of canonical ORFs ribosome-bound and translated are 98.0% and 72%, respectively. Despite this overall similarity, differences were also noted (see “Regulation of translation” below).

For proteomics, we consolidated the results of extensive quantitative proteomics across embryogenesis [26] in our three 8h developmental time windows (methods). This quantitative proteomics data showed an overall 0.45 correlation with our RPKM^FP^ values (Fig. 3C). This correlation appears modest, but it is highly significant (at *p*<0.0001) indicating that RPKM^FP^ can be a good proxy of protein-producing, productive canonical translation. However, 40% canonical ORFs detected as transcribed by RNA-Seq by us and modENCODE during embryogenesis are not detected by proteomics (Fig. 3D), while this number is reduced to 8% by our Poly-Ribo-Seq pipeline. The median RPKM^FP^ value of the proteomics-detected proteins (32.11) is much higher than for those not detected (13.41), indicating that proteomics detection needs high levels of protein translation, as noticed previously [21, 23]. Yet, many ORFs with experimentally-verified translation, and with high Poly-Ribo-Seq metrics, are not detected by proteomics, including all *Hox* genes except the more widely-expressed *Ubx* and *abd-A*. This ‘proteomics detectabilitý issue is not attributable to trypsination treatments, since we observed that 99% of canonical and 92% of non-canonical ORF could produce K- and R-cleaved peptides of 7 to 24aa-long suitable for mass-spec (MS) detection. Hence, other yet unknown factors must influence the proteomics detection of proteins. In addition, small peptides of less than 50 amino acids suffer from enhanced degradation [41, 42], further hindering their detection by proteomics. Indeed, we observed that sCDSs are especially handicapped for MS detection, since the median RPKM^FP^ of those detected is 172.1 (Fig. 3E). Finally, when comparing MS data with RNA-Seq and Poly-Ribo-Seq, and contrary to that observed between RNA and FP data (Fig. 3A) the highest correlation observed is amongst MS data from different embryonic stages. Altogether our analysis suggests that proteomics is an appropriate technique to detect constitutively and highly translated proteins, but not so useful for the detection of ORFs translated either moderately, or in a developmentally-regulated manner.

smORFs also appeared harder to detect by Poly-Ribo-Seq than canonical ORFs, requiring a median RPKM^FP^ of 40.5 (Fig. 2F, 3E). As an independent indicator of translation, we have tagged a sample of smORFs from sCDSs and added to the results from Aspden [23] and [43], for a total of 44 sCDSs. We compared the results of tagging with our Poly-Ribo-Seq pipeline, and with S2 cell proteomics data [44]. Out of the 44 tagged sCDSs, we observed that 43 were detected as translated by tagging, 29 by Poly-Ribo-Seq, and 14 by proteomics (Sup. Fig. 2A). These results showed a coherent pattern, where the three techniques, based on very different detection technologies (Confocal microscopy vs. NextGenSeq vs. Mass-spec, respectively), show decreasing sensitivity thresholds: Tagging > Poly-Ribo-Seq > Proteomics, as also noted when comparing results from other species [45]. Thus, 12 of the 14 proteomics-detected sCDSs were also detected by Poly-Ribo-Seq, and 28 of the 29 detected by Poly-Ribo-Seq were also detected by Tagging. However, our expression of tagged peptides in S2 cells is based on non-endogenous high transcription and this might have pushed the peptide production of lowly expressed ORFs over the edge of detection (see methods and Aspden [23]). Hence, our tagging results may indicate what “can be translated” rather than what actually “is”. Accordingly, 11 of the 13 tag-positive sCDSs with no Poly-Ribo-Seq evidence in S2 cells were translated in other stages. The most abundant tagged peptide localisations (Fig. 3F) were a) reticular-punctate in the cytoplasm, which may correspond to mitochondria [23], or else to mitochondria-associated ER producing the peptide; and b) cortical, which could indicate and association with the plasma membrane or its cytoskeletal cortex. These observations fit with the postulated tendency of sCDSs peptides for membrane-associated localisations [23, 29].

### 4) Regulation of canonical and sCDS translation during embryogenesis

According to our data and modENCODE, 82% of canonical ORFs and 57% of short CDSs were transcribed at any point during development (Fig. 2F). Of these, 8% and 28% respectively were never translated during embryogenesis, giving a first indication of the extent of translational regulation in *Drosophila melanogaster*.

Our data also showed that translation of some 68% of canonical ORFs was detected constitutively (at all embryonic stages), whereas 32% were detected only at specific stages of development (Fig. 4A). Stage-specificity seemed to be higher for smORFs, since only 56% of short CDSs and 18% of uORFs were detected across embryogenesis, whereas 44% and 82% respectively were translated only during some stages of embryogenesis. However, this developmental analysis reflected both transcriptional and translational regulation. To further quantify the extent and dynamics of purely translational regulation, we analysed the changes in translational efficiency (TE) [2] across stages (TE indicates the ratio between FPs and RNA, calculated as RPKM^FP^ / RPKM^RNA^; see methods). This TE analysis revealed both on-off translation changes, and modulations of sustained translation, from one embryonic stage to the next. The average TE varied somehow across development (Fig. 4B), but in general we observed fairly stable and correlated TE values across development (but see also “uORFs as translational regulators”). To identify which ORFs undergo statistically significant translational changes we used the Z-ratio method (which basically indicates whether the variation of a given ORF TE is significantly outside the standard deviation of the total sample; see methods) [46–48]. We also included in the analysis the Unfertilized Egg and S2 cells TE data (comparing them to Early and Late embryogenesis, respectively). We observed significant translational changes in all ORF classes, with the percentage of changed ORFs depending on the ORF class. About 12% of canonical proteins (1,899 of 15,956 analysed), 16% of short CDSs (57 of 355) and 18% of translated uORFs had significantly changed TE from any given stage to the next, indicating significant translational regulation (Fig. 4C). One of the most abundant changes was the translational up-regulation of 539 canonical ORFs from Early to Mid-embryogenesis (Fig. 4D). These ORFs were enriched for GO expression terms indicating a role in organogenesis (i.e. CNS, epidermis and digestive system development) (Fig.4E), as befits the developmental processes underway (Fig. 1A).

**Figure 4.**
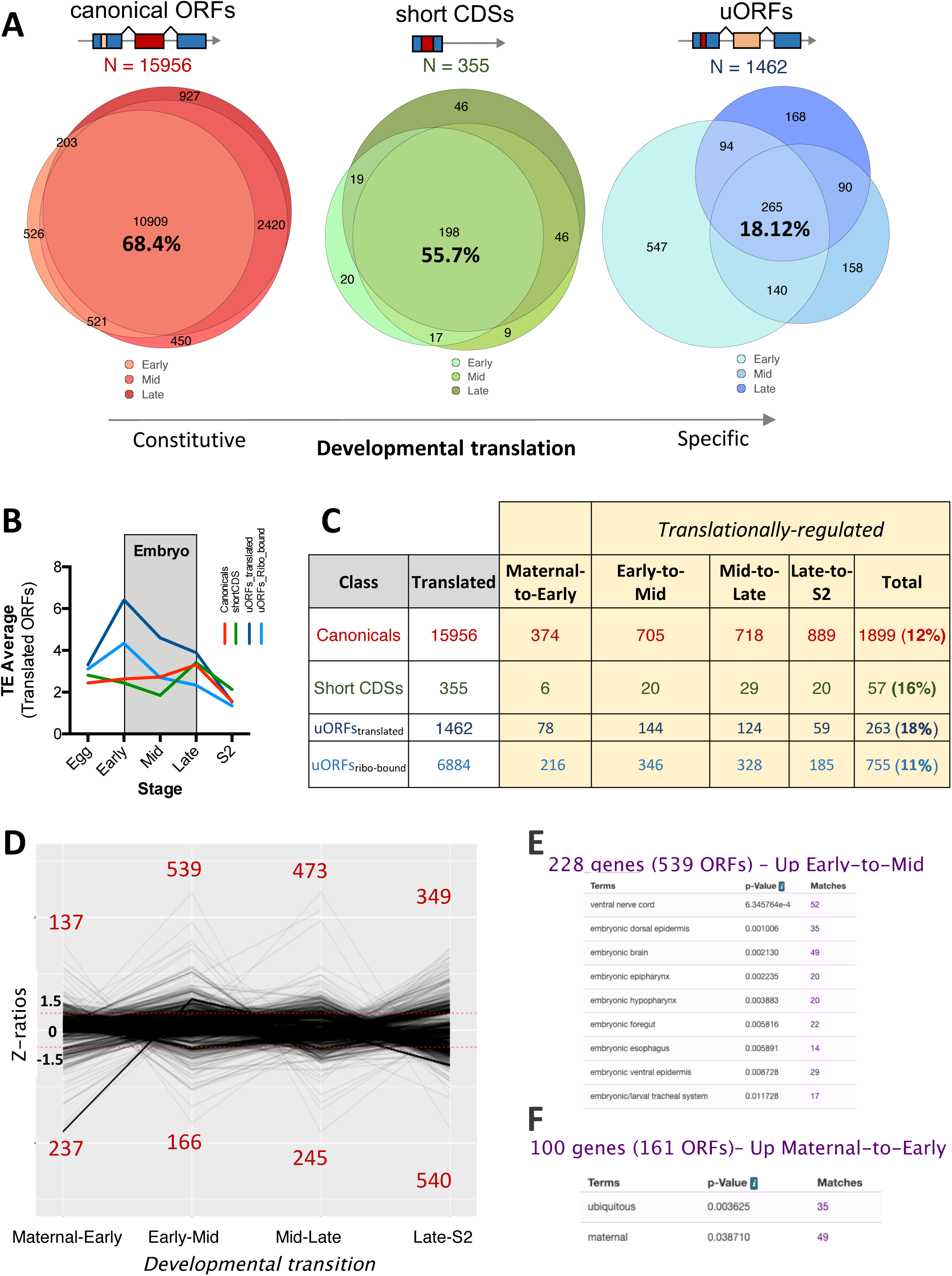
Regulated translation during embryogenesis. **A** Number of transcribed, ribosome-bound and translated ORFs per class across *Drosophila melanogaster* embryonic development, showing progressive specificity in ON/OFF translation patterns. **B** Changes in average translation efficiency, per ORFs class, across analysed developmental time-points. **C** Z-score ratio analysis of translation efficiencies (TE) during successive developmental stages pinpointing ORFs subjected to significant developmental modulations of translation across ORF classes. **D** Number of developmentally-regulated canonical ORFs subjected to downregulation (bottom, Z-ratio ≤ −1.5) or upregulation (top, Z-ratio ≥ 1.5), per developmental transition. **E-F** Significantly-enriched gene expression patterns in the translationally-upregulated canonical genes the Early-to-Mid and Maternal-to-Early developmental transitions respectively (*p-value* of BGDP expression annotations after Benjamini-Hochberg procedure).

The comparison of unfertilized eggs with 0-8h AEL embryo data revealed the maternal to zygotic transition in the translatome. The degradation of maternal mRNAs after the zygotic to maternal transition from 2h AEL, and their substitution by zygotically-transcribed mRNAs has been reported, but a number of mRNAs do persist during early embryogenesis [49]. Our results suggest that some of these maternal mRNAs that persist are subjected to translational repression as an added layer of gene regulation, whereas others go on to increase their translation during embryogenesis. Applying Z-ratio analysis, we observed in total 374 canonical ORFs with significantly changed translational efficiency between the maternal and the early zygotic translatomes (Fig. 4C), of which 237 where down-regulated (including 58 *Dscam* ORFs) and 137 up-regulated, showing RNA expression patterns preferentially registered as ‘maternal’ or ‘ubiquituous’ at this stage (Fig. 4D,F). Further, the maternal to zygotic transition in translation was also highlighted by the correlation between qualitative and quantitative changes in ribosomal binding. 259 maternal-only canonical ORFs appeared translated (framed) in Eggs but not in 0-8h AEL embryos (Fig. 3B), undergoing a significant reduction in ribosomal binding efficiency during this period (as shown by TE, Fig. 3B insert). Reciprocally, 232 ORFs were just ribosome-bound in Oocytes, but fully translated in Early embryos, in correlation with an increase in their TE (Fig. 3B, insert). Finally, 3,170 ORFs were translated in our 0-8h AEL samples but did not yield ribosomal FPs in eggs, presumably corresponding to newly-expressed, zygotic-only ORFs (Fig. 3B).

We hypothesized that a detailed comparison between S2 cells and Late embryos (from where the S2 macrophage-like line was isolated [28]) might reveal the S2 cell-specific translatome profile. Most ORFs seemed translated in both (N=8397), but we observed 382 ORFs translated in S2 cells but not in late embryos, whereas 6,669 ORFs appeared translated in late embryos but not in S2 cells (Sup. Fig. 2B). In addition, 540 ORFs showed significantly lowered TE in S2 cells and 349 significantly raised TE (Fig. 4D). In principle, these differences might reveal a macrophage-like regulatory state, an adaptation to culture, or just a sampling issue (see discussion). Interestingly, we note that the S2 FPs correlate more closely with Early than with Late embryogenesis FPs (Fig. 3A), perhaps reflecting a ‘de-differentiation’ of the S2 macrophage precursors in cell culture.

### 5) uORFs as translational regulators

We obtain highly reproducible Ribo-Seq signal average for uORFs across biological replicates in the Embryo (*r*=0.95, Sup. Fig. 3A), indicating that most uORFs are bound by ribosomes at specific levels across embryogenesis. Indeed, a majority (72%) of embryo-transcribed uORFs show reproducible ribosome binding during embryogenesis (RPKM^FP^>1 in two replicates, Fig. 2F). However, only 21% of these ribosome-binding events show reliable translation signal, as measured by the binomial framing statistic (Fig. 2F), suggesting that a majority of uORFs in *Drosophila melanogaster* might not have a peptide-productive role.

Eukaryotic uORFs can act as cis-translational repressors of a canonical main ORF (mORF) located downstream in their transcripts [50]. This regulatory role has been extrapolated to the transcriptome-wide level as the main function of uORFs [51–54]. Among canonical protein-coding genes, we saw significant shifts in translational regulation across embryogenesis in hundreds of ORFs (Z-ratio ≥ |1.5|) in both Early-to-Mid (705 ORFs) and Mid-to-Late Embryogenesis (718 ORFs; Figure 4B). However a much smaller number of uORFs exhibited significant variation in translation Z-Ratios (144 and 124 uORFs, respectively; Figure 4C), making it unlikely that translated uORFs underpinned most translational changes in canonical ORFs. However, the cis-regulatory role of uORFs is in principle based on ribosome binding and it is mostly independently of the peptide being produced [50, 55, 56], so it could conceivably be carried out by non-productive ribosomal binding too. Therefore, we extended our analysis to include non-framed, ribosome-bound-only uORFs as well (i.e. RPKM^FP^>1 but framing *p>*0.01; Fig. 2A), and this provided a combined pool of 8,346 ribosome-associated uORFs (Fig. 4C).

Previously, a negative correlation between the number of uORFs within a 5’ Leader and the TE of the main ORF was found in Zebrafish genome-wide studies [52] and across vertebrates [53]. We wondered if this was applicable to our 8,346 ribosome-associated uORFs. This would indicate that this negative effect is indeed mediated by uORF ribosomal activity and not a function of other 5’ Leader features [53]. However, we do not see an effect of ribosome-associated uORF number on mORF TE. Instead, we observe a stable TE across mORFs carrying different numbers of uORFs in their 5’Leaders (Fig. 5A).

**Figure 5.**
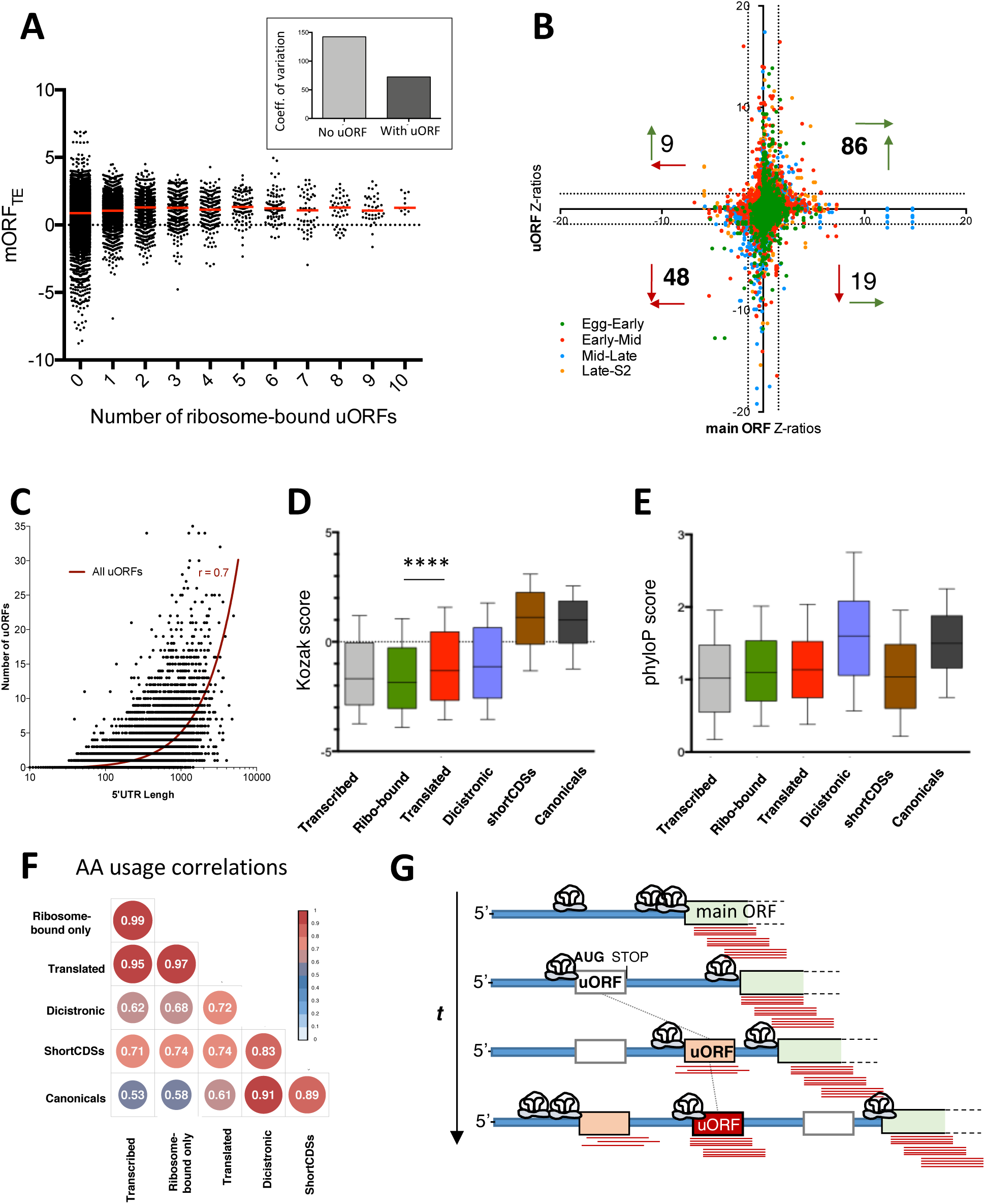
uORFs as translational regulators. **A** Variation in canonical TE levels as a function of the number of cistronic uORFs showing ribosomal binding. **B** Comparison of translational changes between uORFs and cognate canonical ORFs across developmental stages. Numbers denote the number of Z-ratio significant uORF-mORF pairs in each quadrant. Vertical arrows: uORF translational regulation sign. Horizontal arrows: cognate canonical translational regulation sign. Green arrows: Upregulated; Red arrows: Downregulated. **C** Correlation between the number of uORFs per mRNA and 5’UTR length (Spearman’s *r* = 0.7). **D** Comparison of Kozak-context scores between distinct ORF classes and uORF subclasses. Graphs display 10-90 percentile range. Mann-Whitney *p* < 0.001. **E** Comparison of phyloP conservation scores (27-way alignments) between distinct ORF classes and uORF subclasses. Graphs display 10-90 percentile range. Mann-Whitney *p* < 0.01. **F** Correlations in amino acid usage across distinct uORF subclasses, shortCDSs and canonical ORFs; values denote pairwise Spearman’s *r*. **G-H**: FLAG antibody: green, F-actin stained with phalloidin: red. Scale bar denotes 5 micrometres. **G** uORF evolution model. 5’-leaders in blue. t: evolutionary time. Red lines denote ribosomal binding signal; white boxes correspond to novel uORFs; salmon boxes correspond to ribosomal-bound uORFs; red boxes correspond to translated uORFs.

Nonetheless, it would be possible that a minority of uORFs might have important cis-effects on their mORFs. We looked for this ‘uORF onto mORF’ regulation by asking if the 1,037 uORFs with significantly changed TE across stages (270 framed and 767 ribosome-bound-only, Fig. 4C) also show simultaneous significant TE changes in their cis-associated mORF. We found that not to be the case: 79% (817 of 1,037) of uORFs significantly regulated do not have mORFs with significantly changed TE Z-ratios (≥ |1.5|), and vice-versa (Figure 5B). Yet, 12 uORFs correlated negatively with their canonical ORF (i.e. as the uORF TE went up the mORF TE went down, Fig. 5B), suggesting a negative cis-translational regulatory role for these uORFs. Another 25 uORFs displayed reduced TE while their respective mORF TEs were upregulated, which is compatible with such a negative regulatory role. However, 183 uORFs showed coordinated changes with their mORF (i.e. uORF and mORF went both up (124) or both down (59) significantly (Fig. 5B), and this positive correlation extended to the TEs of all 8,504 ribosome-associated uORFs and their mORFs (*r*=0.47 Supp. Fig. 3B). This positive correlation could indicate that uORFs act as either: a) positive translational regulators; b) constant ‘brakes’ or negative regulators that reduce but do not overcome mORF translation; or finally, c) passive ‘bystanderś that are ribosomal-bound by virtue of being present in the 5’Leader of a transcript containing a highly translated mORF. In addition, a combination of these roles is also possible. Interestingly, the total range of TE variation diminishes as the number of ribosome-bound uORFs increases within a given mORF 5’ Leader (Fig. 5A), with canonical ORFs that contain no uORFs in their 5’ Leaders displaying twice the coefficient of variation as those with one or more ribosome-bound uORFs (Fig. 5A, inset). This suggests a cumulative role for uORFs in the maintenance of mORF translational efficiencies, possibly acting as buffers of variation in mORF translation. A buffering effect could also explain the higher TE averages amongst uORFs than amongst canonical ORFs (Fig. 4B), even though the proportion of ORFs with significantly changed TE was identical (1,037 of 8,504 uORFs: 12% vs. 12% canonicals, Fig. 4C - but see also below).

We observed that 45% percent of canonical ORF 5’-Leaders contain uORFs, a number in line with observations across metazoans [25, 29, 57]. Interestingly, this number is significantly lower in shortCDSs (25%). This could be explained by the difference in 5’ Leader average lengths between these two classes (306nt versus 194nt). Indeed, the number of uORFs is positively correlated with the length of the 5’ Leaders across all annotated mRNAs (*r*=0.668, Fig. 5C), indicating that uORFs tend to accumulate more often in longer 5’-Leaders. The positive correlation between uORF number and 5’Leader length supports the idea that uORFs appear randomly in mRNAs [29].

Whether uORFs act as cis-translational regulators, an important number (1,462, Fig. 2F) show framing, i.e. appear to be translated into peptides. On average, RPKM^FP^ (Sup. Fig. 3C) and TE (Fig. 4B) was higher in these framed uORFs, and changed more dramatically across stages than on ribosome-bound-only uORFs (note steeper slope in Fig. 4B and higher total percentage of significant translational regulation in Fig. 4C). The regulation of translated uORF expression was not only quantitative, but included qualitative changes as well (i.e. from ribo-bound-only to framed, correlating with an increase in 32nt FPs mapping to the uORF), and transcriptional regulation, altogether producing that 81% of uORFs were translated in a stage-specific manner (Fig. 4A). Thus, regulated expression seemed a feature of uORF translation.

In the absence of proteomic data to corroborate whether framed uORFs do indeed produce peptides, we checked other bioinformatics markers of coding potential. We observed that translated uORF start codon contexts show significantly better Kozak scores [50] than ribosome-bound-only uORFs (Fig. 5D). Further, the sequence conservation of translated uORFs is intermediate between transcribed uORFs and canonical ORFs (Figure 5E). The amino acid usage produced a similar picture (Figure 5F). These observations might suggest that translated uORFs either produce a specific type of peptide, or that translated uORFs are in an evolutionary transition into coding-ness [29]. One end result of such a transition would be to produce a di-cistronic coding gene (uORF plus mORF), and it is interesting that genes annotated as such by FlyBase display similar conservation to canonical ORFs (Fig. 5E), but aminoacid usage intermediate between these and translated uORFs (Fig. 5F).

## DISCUSSION

### Translation vs. peptide function

Protein translation is the most expensive gene activity undertaken by a cell, one order of magnitude more costly than either replication or transcription, and costly enough to have a selective impact [58]. However, it is important to disentangle the act of productive translation from its biological purpose. It is usually expected that translated ORFs must produce a peptide or a protein with canonical function. However, a translated peptide could have other biological activities, [59] or no peptide function at all, such as the act of translation could be the biologically relevant function of the encoding ORF (as in regulatory uORFs, or in canonical ORFs during nonsense-mediated decay) [55, 60]. Ribo-Seq, and other techniques such as proteomics, do not reveal protein and peptide function, only its expression. Protein function is suggested by other studies (such as sequence conservation) and ultimately tested experimentally.

smORF translation is difficult to ascertain, but has been repeatedly detected to occur at a large scale by Ribo-Seq, including in this study. Different types of studies and analyses yield numbers always in the thousands per genome [45]. These numbers are a small fraction of the smORFs in the genome (about 5%), yet it is large enough (about a 10% of extra coding genes to add to canonical genes) to present a challenge to our understanding of genomes and their function. What can be the biological purpose of this 10% ‘crypto-translatome’? Due to their shortness, it is challenging to determine smORF function from their sequence. GO analysis of the longer, and annotated, sCDS smORF class, and analyses of their aminoacid composition [23, 29], suggested roles related to cell membranes and organelles. However, shorter smORFs such as uORFs and those found in lncRNAs (called lncORFs) are not annotated and do not display known protein domains, nor homologies to characterised proteins, that could suggest a peptide function. Their AA composition is intermediate, yet distinct from canonical proteins and random sequences [29]. Nonetheless, important functions for lncORF and uORF peptides have been proven experimentally in a few cases [35, 39, 61–64], as well as a cis-regulatory, ‘non-coding’ function for uORFs [50] [55, 56].

### Ribosomal binding versus productive translation

Our results suggest a distinction between ribosomal-binding-only and productive translation. We observed that about 59% of transcribed uORFs show abundant, yet not framed, ribosomal binding. This is unlike canonical ORFs and short CDSs, where only 5% and 18% respectively show ribosomal association not corresponding to productive translation. This raises the possibility that either much of ribosomal binding in uORFs serve purposes other than peptide production, or that it reflects ‘translational noise’. In view of the energetic and selective costs involved, it is unlikely for the cells to allow for their resources to just be ‘lost in translation’. An explanation might be suggested by the maternal translatome.

### Maternal Translation

The maternal to zygotic transition corroborates that ‘ribosomal-binding-only’ without framing is a different state than fully-fledged translation. Our data revealed ORFs ‘poised’ for translation between unfertilized eggs and Early embryos. 232 canonical ORFs switch from unproductive ribosome-bound-only to productive translation, while increasing their translational efficiency (Fig. 3B). In other words, the quality of their ribosomal association changes from low affinity (low TE) and not following a particular frame, to higher-affinity, frame-linked productive translation. To produce non-framing, ribosomal binding must happen in overlapping frames, or ‘shift’ between frames, or include ‘scanning’ 40S subunits, or proofreading ribosomal units involved in NMD and not productively reading codons. Interestingly, we observe one-frame shift of ribosomes at STOP codons (from blue frame “2” to red “0”, Fig. 2B,E), also noticeable in other Ribo-Seq studies [21], which may reflect a conformational change or the ribosome pausing for a time at its A site while disengaging from the mRNA. Under weak translation conditions (indicated by lower RPKM^FP^, Sup. Fig. 3C), in some ORFs elongation may proceed more slowly (perhaps due to a failure to acquire AA-loaded tRNAs or elongation factors). It has been shown that prolonged ribosomal stalling plus queuing can produce both premature termination [65], and use of non-AUG START codons in a different frame [66]. Either effect could produce a frameshifted Ribo-Seq signal towards frames “0” and “1” as we observe, distorting the main ORF signal in frame “2”. In other words, ribosome-bound-only ORFs could be failing to produce enough START to STOP translation, while increasing a frameshifted, noisy ribosomal binding. This effect could be temporary (as in the maternal to zygotic transition) or a constitutive feature of the ORF (as in most uORFs, see below).

### Function of uORFs

In our previous study [23] we estimated that 34% of Drosophila uORFs were translated in S2 cells, on the basis of a high RPKM^FP^. Interestingly, our improved methods reveal two uORF pools, whose RPKM^FP^ values overlap yet having different averages, and different framing (Sup. Fig. 3C). The percentage of ribosome-bound uORFs in *Drosophila* embryos (71%), is similar to recent estimates [54], but the percentage of actually translated uORFs in our samples is only 12%. The significance of non-productive ribosomal-binding-only in 59% of transcribed uORFs is intriguing. The positive correlation between uORF and mORF with significantly changed TEs, the genome-wide lack of effect of ribosome-bound uORFs on mORF TE, and the correlation between 5’Leader length and number of uORFs all seem to suggest that most uORF are a random and function-less by-product of 5’Leader sequence variation, also suggested by the random-like size distribution of uORFs [29]. Yet, this does not exclude that some uORFs could have a cis-regulatory, or a peptide-producing role. Different mechanisms have been proposed to mediate a cis-regulatory function, including ribosome stalling and disengagement at uORFs, a reduction in translational re-initiation on the downstream mORF AUG codons, and the triggering of mRNA decay [50, 56, 67], all leading to a decrease in mORF translational efficiency. Positive regulatory roles are also possible, with uORFs recruiting ribosomes and thus increasing the likelihood that they reach mORFs via reinitiation, and/or imposing an appropriate ribosomal spacing (reducing ribosome collisions and thus improving mORF TE and fidelity [65, 66, 68]). We propose a combined role, with most uORFs acting as ‘translation buffers’ that would recruit ribosomes (a relatively scarce yet valuable cellular resource [68, 69]) and pass them on to their mORF at a constant rate, while discarding ribosomes in excess. A passive, low-level uORF buffer effect at the genome-wide level is consistent with studies in yeasts [70, 71] vertebrates [25] and humans where a genome-wide positive correlation between the TE of individual uORFs and their cognate mORFs is also observed. A function of uORFs as buffers could also generate stalling, queuing, termination and frameshifted starts [66], and explain their frameshifted FP signal.

A mildly beneficial buffering role could explain the ubiquity of uORFs in otherwise canonical mRNAs. The minority of uORFs showing peptide-productive translation could be a tolerable by-product of this buffering role, which would otherwise be mostly mediated by the much more prevalent uORFs with non-productive ribosomal-binding-only. Interestingly, we observe subtly different Kozak scores, conservation levels and aminoacid usage for translated uORFs, which seem to fit into a stepped continuum ranging from transcribed-only uORFs to canonical ORFs. It is tempting to speculate that, just as ribosomal-bound-only in canonical ORFs may be a developmental transitory state during the maternal to zygotic translation, ribosome-bound-only uORFs may be an evolutionary transitory state poised for a transition to full coding function. In this way, the 5’Leaders of canonical genes would act as ‘proto-gene nurseries’ offering a tolerant environment for random ORF creation, while providing coding mRNA features such as poly-A tails, splicing-related and translationally-related RNA processing and stability, allocation to polysomes, and a specific transcriptional pattern. Dicistronic genes could be another step in this transition from inert or regulatory uORF to coding ORF (although dicistronics can also emerge from the fusion of two pre-existing canonical ORFs). Interestingly, short CDSs, which in most aspects studied here (RPKM reproducibility; translation percentage; MassSpec detection; embryonic stage specificity; prevalence of translation regulation; amino-acid usage) are intermediate between translated uORFs and canonical ORFs, do not show intermediate conservation (Fig. 5E). The lower sCDSs conservation may suggest that sCDSs are also involved in another evolutionary process, perhaps another and more dynamic stepped continuum leading to coding-ness via lncRNAs [29, 72].

### Further developments-single cell translatomics

Despite the improvements to our Poly-Ribo-Seq protocol, and the large number of reads that we have collected, it is possible that not all embryonic translation has been detected in our samples. Lowly abundant or transient mRNAs may not be adequately detected in our 8-hour developmental windows, or may be drowned out by noise and not be able to achieve RPKM >1. This could be especially true of mRNAs expressed in a few cells of the embryo, a situation that must arise often towards the end of embryogenesis, and could be prevalent in differentiated tissues. Our macrophage-like S2 cell line data allows for an approximation to this issue, and the problem of integrating and comparing Ribo-Seq and genomic data from whole embryos and cell lines. Direct comparison between late embryos and S2 cells creates a sampling issue. The embryonic counterparts of S2 cells could be only a few cells amongst many embryonic cell types and some 30,000 cells overall [73, 74]. Thus, the FPs from S2-related ORFs might not be adequately detected in whole embryo extractions (thus yielding a low or no RPKM), whereas the same FPs will be well-detected in the monotypic S2 cell culture. For example, we showed the specific expression (by *in situ* hybridisation and RT-PCR) and function (by observation of mutant phenotypes in embryonic macrophages) of the *hemotin* smORF gene in the *Drosophila* embryo [75]. Yet our RNA-Seq and Ribo-Seq data, and modENCODE RNA-Seq would indicate that *hemotin* is neither transcribed nor translated during embryogenesis, while the same data show vigorous expression in S2 cells (RPKM^FP^= 58.6 and framing *p*=8.15 x e10^-5^). This sampling issue can contrive an artefactual increase of expression and translation (RPKM and TE) of *hemotin* and similar cell-specific transcripts in cell lines [76] when compared with whole embryos or organs. However, a lowering of RPKM or TE in S2 cell cultures in 541 ORFs (Fig. 4D) cannot be due to this sampling issue, and must reflect specific translational repression in a S2 ‘cell fate’, following either a macrophage-like fate, or a cell-in-culture fate. Thus, Ribo-Seq data from cell lines can reveal genes whose translation declines during specific cell differentiation programmes. Ideally, it would be best to take RiboSeq to this single cell level. The demand for high levels of starting material (in the region of µg) is the largest barrier to single-cell Ribo-Seq, but it could be overcome by the combination of Ribosomal Immunoprecipitation (TRAP) (which requires even more starting material, in the region of mg [77, 78]), with further improvements to the core polysome purification protocol [79].

## CONCLUSIONS

The likelihood of translation can be determined in Ribo-Seq data by comparing the frequency of reads in frame against a simple binomial probability. Using this metric and a standard RPKM>1 filter for ribosomal binding, almost 16,000 canonical ORFs, more than 300 sCDSs and almost 1,500 uORFs appear translated during the embryonic development of *Drosophila melanogaster.* Translation levels appear more reproducible, yet highly correlated, with transcription, together yielding highly constant expression of canonical ORFs and sCDSs, and higher temporal specificity of uORFs. However, 12-18% ORFs show specific regulation of their translation, including ‘poising’ at the maternal to zygotic transition. Finally, a large pool of near 7,000 uORFs also show ribosomal binding without achieving either productive translation, or a significant regulatory input on downstream ORFs. These and other data suggest that, in general, 5’-Leaders with uORFs act as ‘translation buffers’ at the gene level (helping downstream ORFs to stabilize translation levels), and as ‘proto-gene nurseries’ at the genomic level (providing a favourable environment for random ORF creation).

## MATERIALS AND METHODS

### Ribo-Seq and RNA-seq procedures

The RNA-Seq and Poly-Ribo-Seq experiments were conducted as previously described in Aspden et al. [23] with the following modifications. Embryos were flash frozen, turned into powder using a pre-chilled pestle and mortar and homogenized in Lysis Buffer at 4C for 20 min. Pre-clarified lysates were processed for mRNA isolation and fragmentation and polysome separation and digestion as previously described. Digested polysome samples were concentrated by centrifugation and subsequently loaded into a 1M Sucrose Cushion and centrifuged at 70,000g to pellet the monosomes. The 50nt fragmented mRNAs and 28-34 nt footprints were isolated from 10% denaturing acrylamide gels and subsequently T4 PNK treated for library preparation.

The NEBNext Small RNA Library Prep Set for Illumina (NEB, E7300) was used following manufacturers’ instructions with small modifications as follows. After the 3’Adapters ligation step, ribosome footprints were rRNA depleted using biotinylated DNA fragments and oligos from rRNAs and going through two consecutive rounds of subtractive hybridisation (except for the T 0-8h sample having only one round) using MyOne Streptavidin C1 Dynabeads (Invitrogen). After Ethanol precipitation, samples were processed as NEB guidelines. Libraries were isolated from 10% non-denaturing gels by size selection and their quality and quantity measure by DNA high sensitivity chips (Agilent) with Bioanalyzer and Qubit.

### Footprint sequence alignment

Ribosomal Footprints were filtered (PHRED quality≥33) and clipped for adapters using the FASTX-Toolkit. The resulting reads were aligned to a FASTA file containing rRNA, tRNA, snoRNA and snRNA from Flybase Release 6.13 annotations using Bowtie, discarding the successful alignments and collecting all unaligned reads. Unaligned reads were then mapped to the FlyBase *D.melanogaster* reference genome (Release 6.13) using HISAT2 with default options. Due to the very low frequency of multimapping reads, we retained all genome-aligned reads for further analyses.

### ORF selection and identification

All annotated coding sequences (CDSs) in FlyBase, r6.13 were retrieved and divided into groups of either longer than or shorter than 303 nucleotides (100 AA). CDSs longer than the cut off were assigned to the canonical ORF category. CDSs shorter or equal to 303 nt. but arising from the same locus as canonicals (short-isoform smORFs, [29]) were discarded from our analysis. The remaining short CDSs, which arise from independent genomic loci, were assigned to the short CDS category [29]. We identified uORFs longer than 10 aa with an AUG start codon followed by an in-frame stop codon within the annotated 5′ Leaders of all transcripts annotated as protein-coding in FlyBase r6.13, using the emboss *getorf* program. Next, all ORFs with fully-redundant CDS coordinates were identified, and duplicates were discarded within each class (canonicals, short CDSs and uORFs), Additionally, we only kept uORFs with no overlap with any annotated CDS. uORFs in dicistronic transcripts were then added to the uORF set, based on a full overlap with annotated ORFs contained within the 5’ Leader of an immediately downstream annotated CDS.

### Ribosomal Footprint analysis

Relative Ribosome density (RPKM^FP^) was measured by counting Ribo-Seq reads overlapping each feature using the R package Rsamtols (absolute ribosome density), and scaling this number by feature length and the total number of genome-aligned reads [2]. Total mRNA expression (RPKM^RNA^) was ascertained using the same features and applying the same method on RNA-Seq measurements. Correlations analyses across replicates were measured by Spearman’s rho on ORF RPKM values.

### Detection of framing and individual translation events

To analyse framing, ribosome footprints (FPs) were first aligned to an artificially constructed transcriptome set, consisting of all ORFs analysed in this study as well as their surrounding regions −18 nucleotides upstream and +15 nucleotides downstream, thus allowing for the alignment of full ribosomal reads spanning the START and STOP codons. The resulting alignments were analysed using the R package RiboSeqR to extract the global framing patterns across ribosome footprint lengths for three pools of RPFs: all pooled embryonic RiboSeq samples, S2 cell samples and pool of mature oocyte and activated eggs RPFs (Kronja et al. [18], cicloheximide^+^). The main read length and its most overrepresented frame in each sample was then used to evaluate the framing of each ORF in each Ribo-Seq sample. For each ORF, we then compared successful framing events (those matching global framing patterns) with unsuccessful framing events using the binomial test implemented *pbinom()* function in R to measure the probability of obtaining the observed framing (or better), when compared to the expectation by random (1 in 3 probability). For binomial tests of ORF translation, different biological replicates (if available) were merged into a single one. Although the precision of the binomial probability method improves with the amount of evidence (i.e. higher number of reads can achieve very low p-values, or very high probabilities of being translated, Fig. 2C), it can still achieve results with minimal information, which is especially relevant for smORFs as their short size accrues less reads in sequencing experiments when compared to canonical ORFs with the same RPKM range (Fig. 1, 2E). Thus, a minimum of 5 reads is required to obtain p<0.01 if all reads are in frame, or more than 7 reads if any is out of frame [38].

### Comparison with other Ribo-Seq and RNA-Seq datasets

We obtained previously-published [18] fastq files containing Cycloheximide^+^ Ribo-Seq reads from mature oocytes and activated eggs from the NCBI Gene Expression Omnibus (accessions SRR1039770 and SRR1039771) as well as the corresponding RNA-Seq experiments (SRR1039764 and SRR1039765), all deposited under accession number GSE52799. The Activated Egg and Oocyte reads were merged to obtain Ribo-Seq and RNA-Seq “Unfertilized Egg” samples, which were analysed employing the same pipeline as our own data. In the case of S2-cell data, we merged our previously-published Ribo-Seq (accessions SRR1548656, SRR1548657, SRR1548658, SRR1548659) and RNA-Seq datasets (SRR1548660-SRR1548661) [23]). We obtained modENCODE RNA-Seq 2-hourly embryonic-staged datasets from FlyBase and consolidated them into our 8-hour-long stages (0h-8h, 8h-16h and 16h-24h AEL).

### Comparison between RNA-seq and mass spectrometry datasets

We compared the mass spectrometry-based developmental proteome of *Drosophila melanogaster* from [26] with our own sequencing data by first averaging the imputed log2 LFQ intensity per protein-group across samples spanning embryonic windows matching those of our timed collections (0h-8h, 8h-16h and 16h-24h AEL). Finally, we normalized all datasets and measured the correlation between the average protein intensities for all identified proteins and our own gene expression measurements using Spearman’s rho.

### Translation Efficiency and Translational Regulation analysis

Translation efficiency (TE) per ORF was calculated as the fraction of ribosome-density (Poly-Ribo-Seq RPKM) over the total mRNA read density (RNA-Seq RPKM). To detect significant events of translational regulation, we performed Z-ratio calculations of TE variation as per Cheadle et al. 2013 [48]. First, TE values were normalized by calculating the Z-score of each ORF in each sample:

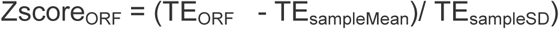

Second, Z-ratios across contiguous developmental stages e.g. Early and Mid-embryogenesis were calculated for each ORF:

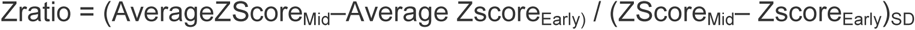

Third, the cut-off of ≥|1.5|was used to identify biologically significant differences [47, 48] between the two contiguous stages.

### Gene Expression enrichment analysis

We used the unique gene identifiers (FBgn) for genes identified as translationally-regulated across stages (see Translational Regulation analysis) to calculate the embryonic tissue-expression enrichment using the Intermine tool (Flymine), which uses ImaGO terms associated with expression patterns deposited in the BDGP database. The enrichment for each gene set was measured against a background of all *Drosophila* melanogaster genes with expression information, and expressed as a p-value for each significantly-enriched tissue, after the Benjamini-Hochberg multiple-comparisons correction.

### Conservation analysis

The pre-computed phyloP nucleotide scores for 27-way insect multiple sequence alignments were downloaded from UCSC as bigWig files. We then used the UCSC bigWigAverageOverBed package to compare phyloP scores with BED files containing genome-coordinates for each ORF in our set, obtaining average phyloP scores per ORF.

### Amino acid composition analysis

We used a previously published own script (*aa_composition.pl*, [23]) to calculate the observed or predicted amino acid compositions of our different sets of ORFs.

### Kozak-context scoring per ORF

For all canonical ORFs, we extracted the nucleotide composition around-but excluding- the annotated START codon (−5 to 6, excluding positions 1-3). For each position, we then calculated the log odds ratio between observed and background nucleotide frequencies. This provided a scoring table of position-specific nucleotide frequencies from *bona-fide* Kozak contexts with which to score individual ORFs in all classes. The final score per ORF was then obtained by adding the individual position-specific values for all observed nucleotides.

## DECLARATIONS

### Ethics approval and consent to participate

Not applicable

### Consent for publication

Not applicable

### Availability of data and materials

Data is retained by the corresponding author and will be made available in a public repository upon publication

### Authors’ contributions

PP: conception and overall design of the work, bioinformatics analysis and interpretation of Poly-Ribo-Seq, proteomics and genomics data; writing of software and drafting the manuscript. MASM: design of Poly-Ribo-Seq experiments and acquisition of Poly-Ribo-Seq data. JIP: acquisition, bioinformatics analysis and interpretation of Poly-Ribo-Seq data; design and acquisition of tagging data; substantial revision of the manuscript. JLA: interpretation of Poly-Ribo-Seq data; design and acquisition of tagging data, substantial revision of the manuscript. JPC: conception and overall design of the work, interpretation of tagging and Poly-Ribo-Seq data, drafted the manuscript.

## Acknowledgements

We thank Rose Phillips for excellent technical assistance, Unum Amin for transfection and confocal microscopy experiments, and Emile Magny and Sarah Newbury for discussions and comments to the manuscript. We also thank Terry Orr-Weaver and Stephen Eichhorn for providing information on the Kronja et al. data.

---

**Supplementary Figure 1 – RiboSeq read length distribution and transcriptome-wide framing. A** Distribution of Poly-Ribo-Seq genome-mapped ribosome-protected fragment (RPFs) lengths; codon-framing of predominant lengths shown in inset. **B** negative control: of codon-framing in genome-mapped RNA-Seq reads fails to show dominance in any of the frames.

**Supplementary Figure 2 –A** Detectability of mRNA translation across techniques. Total number of sCDSs detected by Mass-spectrometry, Poly-Ribo-Seq and FLAG-Tagging (Total N=44). **B** S2-cell canonical translation. Comparison of translated canonical ORFs in S2 cells and the late embryo stages from which these cell culture primarily originated [28].

**Supplementary Figure 3 – uORF A** Comparison of uORF translation from T and B replicas (in log_2_RPKM) show a very high correlation (Spearman’s *r*=0.9466). **B** Histogram of individual Spearman’s correlation *r* between TE of ribosome-bound uORFs and their corresponding mORF across embryogenesis. The overall positive correlation has a median of *r=*0.477 (binning=0.1 windows). **C**. Distribution of log_2_(RPKM^FP^) values of ribosome-bound-only uORFs (red) and translated (framed) uORFs (green).

**Supplementary Table 1 – Sequencing statistics for novel RiboSeq and RNA-Seq datasets reported in this study.** Number of raw reads per replica per stage and percentage of remaining reads after ribosomal RNA subtraction (“norRNA”) and genome-mapping (see methods).

**Supplementary Table 2 – Heatmap of correlations across RNA-Seq and RiboSeq samples and replicates** Spearman correlation values (*r*) for canonical ORF RPKMs across all RNA-Seq and RiboSeq replicates analysed in this study. All correlations were significant (p<0.0001). T and B denote the two biological replicates used in this study. Early, Mid and Late denote Embryonic windows 0-8h, 8-16h and 16-24h, respectively. “Flybase” RNA-Seq refers to modENCODE data (Graveley et al. 2011). S2 refers to S2-cell datasets (see methods). Unfertilized Egg corresponds to previously-published data (Kronja et al. 2014).

**Supplementary Tables 3 - Gene expression, regulation and conservation statistics for distinct small ORFs in polycistronic mRNAs. A** The *tal* polycistronic mRNA shows marked quantitative differences in translation efficiency, probability and regulation, as well as conservation, across its four small ORFs. **B** The *scl* locus contains two cistronic small ORFs that show similar but quantitatively different levels of translation and conservation.

**Supplementary Table 4 – Number of Transcribed, Ribosome-Bound and Translated ORFs by Class and embryonic stage.** Different ORF classes show distinct amounts of transcription, ribosome binding and translation across stages.

