## Supplemental Figures and tables for "Developmental regulation of Canonical and small ORF translation from mRNAs"

### Supplementary Figure 1

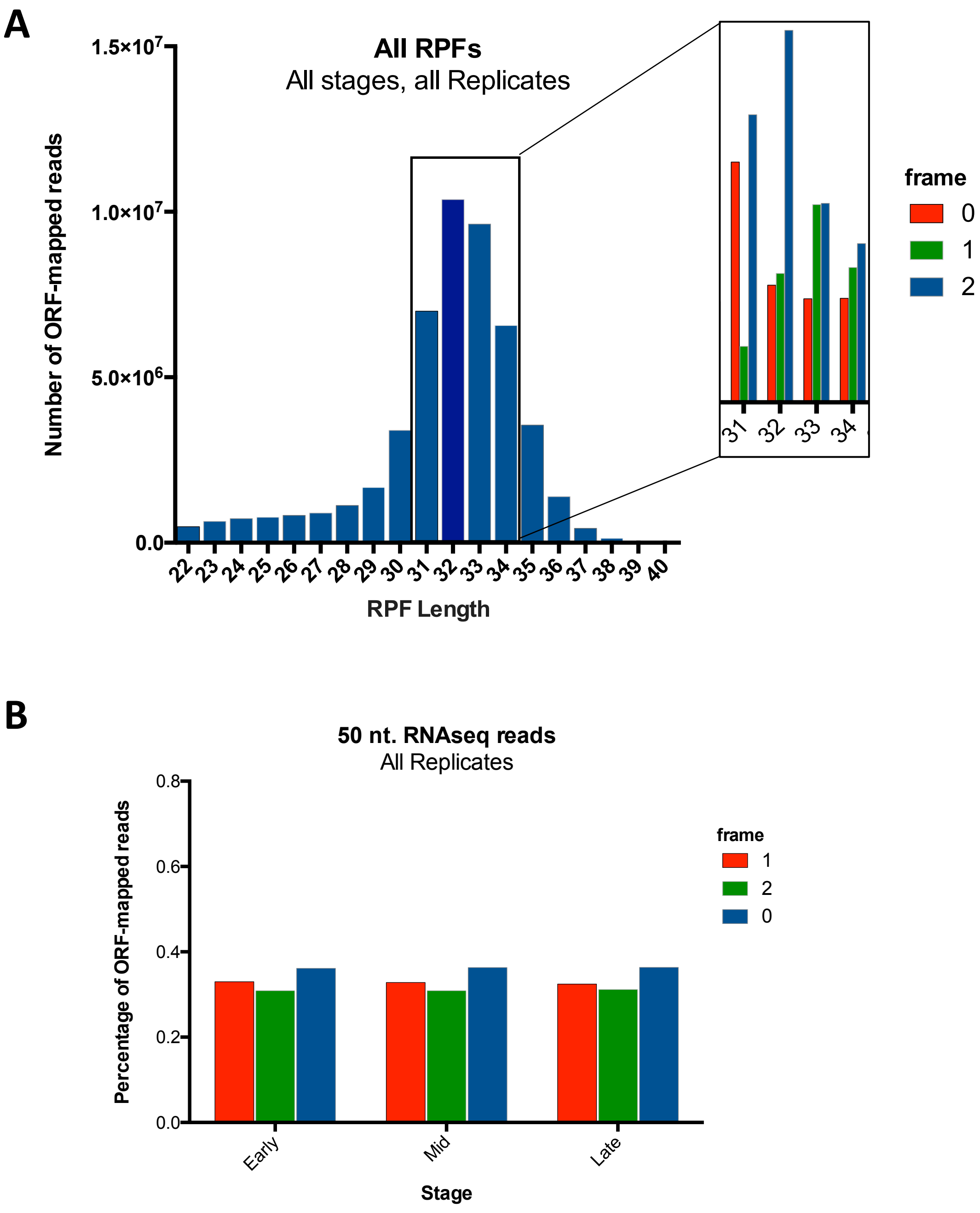

#### Supplementary Figure 2

**A**

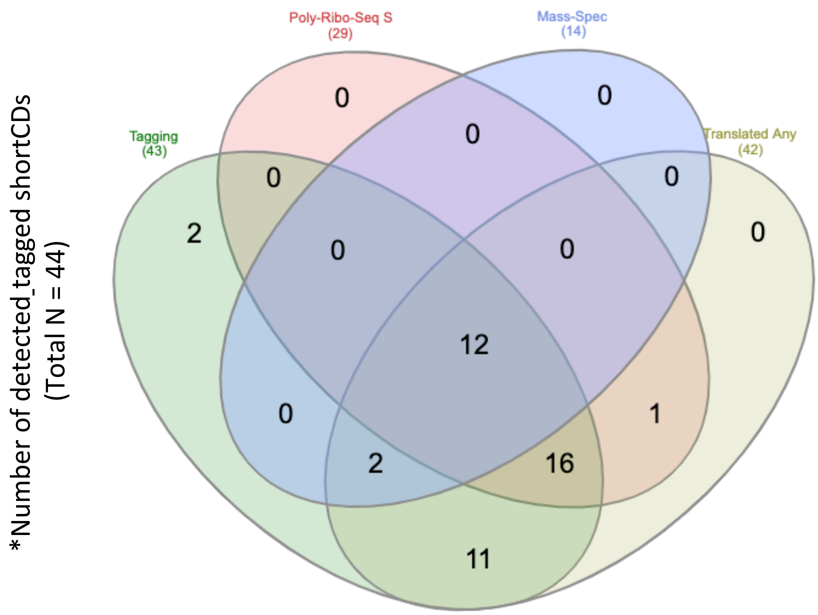


**B**

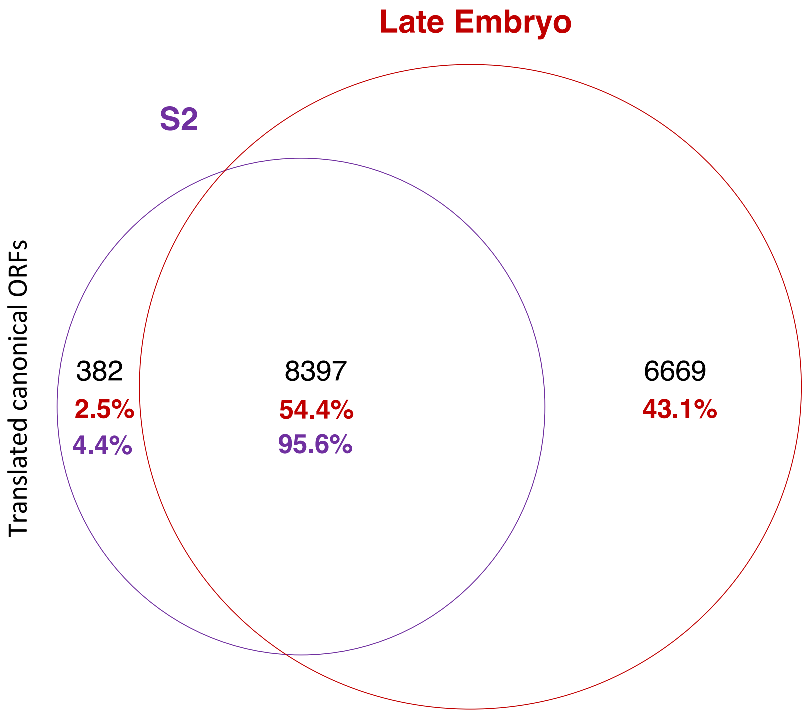

### Supplementary Figure 3

A

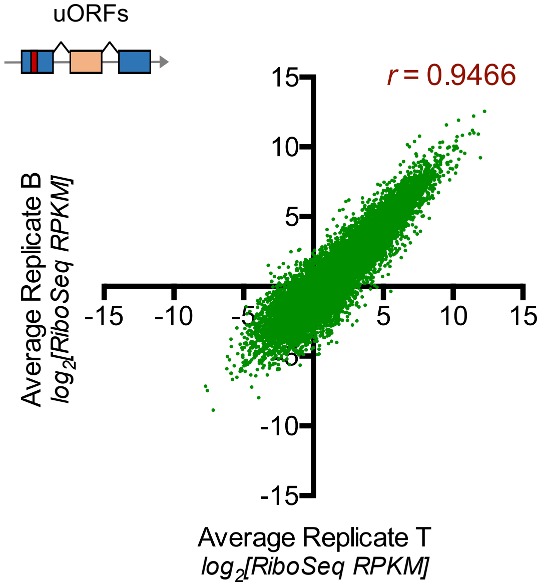


B

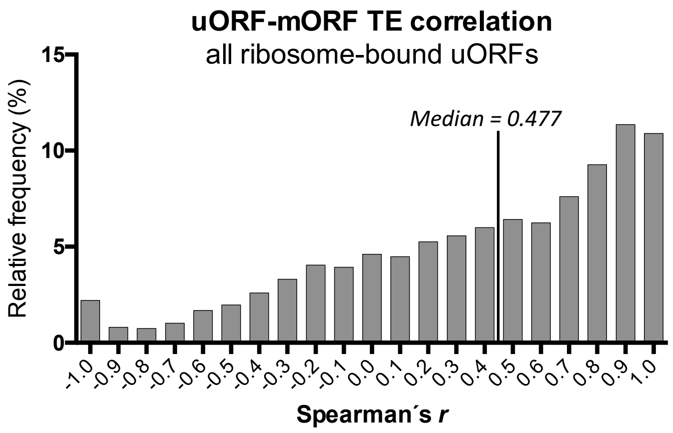


C

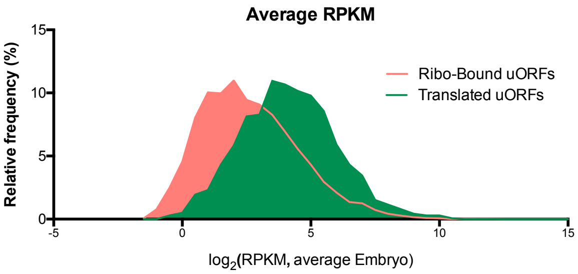

### Supplementary Table 1

| | | | Reads | | |
|---|---|---|---|---|---|
| Sequencing | Replica | Stage | Raw reads | norRNA (%) | Genome-mapped (%) |
| RiboSeq | T | Early | 285.529.728 | 14.8 | 73 |
| | B | Early | 322.959.034 | 41.6 | 74.5 |
| | T | Mid | 178.496.111 | 41.2 | 75.2 |
| | B | Mid | 170.312.394 | 29.3 | 82.1 |
| | T | Late | 110.156.905 | 72.4 | 77.2 |
| | B | Late | 158.351.452 | 47.2 | 75 |
| RNA-Seq | T | Early | 39.817.649 | 51.8 | 74.8 |
| | B | Early | 99.986.496 | 87.6 | 84.5 |
| | T | Mid | 54.333.403 | 44.5 | 56 |
| | B | Mid | 170.312.394 | 49.8 | 79.8 |
| | T | Late | 68.857.596 | 53.5 | 53.9 |
| | B | Late | 66.245.016 | 74.7 | 71.9 |

### Supplementary Table 2

| | Early_RNA-Seq_Flybase | Mid_RNA-Seq_Flybase | Late_RNA-Seq_Flybase | Early_Riboseq_T | Early_Riboseq_B | Mid_Riboseq_T | Mid_Riboseq_B | Late_Riboseq_T | Late_Riboseq_B | Early_RNA-Seq_T | Early_RNA-Seq_B | Mid_RNA-Seq_T | Mid_RNA-Seq_B | Late_RNA-Seq_T | Late_RNA-Seq_B | S2_Riboseq | S2_RNA-Seq | Unfertilized_Egg_Riboseq | Unfertilized_Egg_RNA-Seq |
|---|---|---|---|---|---|---|---|---|---|---|---|---|---|---|---|---|---|---|---|
| Early_RNA-Seq_Flybase | 0 | 0.87 | 0.64 | 0.93 | 0.92 | 0.68 | 0.71 | 0.53 | 0.53 | 0.83 | 0.94 | 0.66 | 0.69 | 0.54 | 0.62 | 0.81 | 0.8 | 0.86 | 0.86 |
| Mid_RNA-Seq_Flybase | 0.87 | 0 | 0.82 | 0.84 | 0.85 | 0.86 | 0.87 | 0.68 | 0.67 | 0.76 | 0.84 | 0.82 | 0.84 | 0.7 | 0.71 | 0.76 | 0.74 | 0.75 | 0.72 |
| Late_RNA-Seq_Flybase | 0.64 | 0.82 | 0 | 0.65 | 0.67 | 0.87 | 0.82 | 0.89 | 0.86 | 0.65 | 0.65 | 0.82 | 0.83 | 0.85 | 0.83 | 0.61 | 0.58 | 0.57 | 0.54 |
| Early_Riboseq_T | 0.93 | 0.84 | 0.65 | 0 | 0.99 | 0.75 | 0.79 | 0.61 | 0.63 | 0.85 | 0.95 | 0.72 | 0.73 | 0.58 | 0.69 | 0.8 | 0.78 | 0.88 | 0.83 |
| Early_Riboseq_B | 0.92 | 0.85 | 0.67 | 0.99 | 0 | 0.78 | 0.82 | 0.64 | 0.66 | 0.86 | 0.95 | 0.74 | 0.76 | 0.61 | 0.72 | 0.79 | 0.77 | 0.86 | 0.81 |
| Mid_Riboseq_T | 0.68 | 0.86 | 0.87 | 0.75 | 0.78 | 0 | 0.98 | 0.88 | 0.87 | 0.71 | 0.73 | 0.91 | 0.9 | 0.82 | 0.83 | 0.66 | 0.63 | 0.63 | 0.59 |
| Mid_Riboseq_B | 0.71 | 0.87 | 0.82 | 0.79 | 0.82 | 0.98 | 0 | 0.84 | 0.85 | 0.74 | 0.76 | 0.9 | 0.91 | 0.79 | 0.82 | 0.69 | 0.66 | 0.67 | 0.62 |
| Late_Riboseq_T | 0.53 | 0.68 | 0.89 | 0.61 | 0.64 | 0.88 | 0.84 | 0 | 0.99 | 0.64 | 0.61 | 0.82 | 0.82 | 0.88 | 0.89 | 0.57 | 0.54 | 0.52 | 0.48 |
| Late_Riboseq_B | 0.53 | 0.67 | 0.86 | 0.63 | 0.66 | 0.87 | 0.85 | 0.99 | | 0.65 | 0.62 | 0.81 | 0.81 | 0.86 | 0.89 | 0.58 | 0.55 | 0.54 | 0.5 |
| Early_RNA-Seq_T | 0.83 | 0.76 | 0.65 | 0.85 | 0.86 | 0.71 | 0.74 | 0.64 | 0.65 | 0 | 0.92 | 0.83 | 0.83 | 0.72 | 0.82 | 0.74 | 0.75 | 0.76 | 0.79 |
| Early_RNA-Seq_B | 0.94 | 0.84 | 0.65 | 0.95 | 0.95 | 0.73 | 0.76 | 0.61 | 0.62 | 0.92 | 0 | 0.75 | 0.79 | 0.64 | 0.73 | 0.82 | 0.81 | 0.87 | 0.88 |
| Mid_RNA-Seq_T | 0.66 | 0.82 | 0.82 | 0.72 | 0.74 | 0.91 | 0.9 | 0.82 | 0.81 | 0.83 | 0.75 | 0 | 0.95 | 0.88 | 0.91 | 0.64 | 0.63 | 0.61 | 0.6 |
| Mid_RNA-Seq_B | 0.69 | 0.84 | 0.83 | 0.73 | 0.76 | 0.9 | 0.91 | 0.82 | 0.81 | 0.83 | 0.79 | 0.95 | 0 | 0.89 | 0.89 | 0.68 | 0.68 | 0.63 | 0.62 |
| Late_RNA-Seq_T | 0.54 | 0.7 | 0.85 | 0.58 | 0.61 | 0.82 | 0.79 | 0.88 | 0.86 | 0.72 | 0.64 | 0.88 | 0.89 | 0 | 0.91 | 0.57 | 0.56 | 0.5 | 0.49 |
| Late_RNA-Seq_B | 0.62 | 0.71 | 0.83 | 0.69 | 0.72 | 0.83 | 0.82 | 0.89 | 0.89 | 0.82 | 0.73 | 0.91 | 0.89 | 0.91 | 0 | 0.61 | 0.6 | 0.59 | 0.57 |
| S2_Riboseq | 0.81 | 0.76 | 0.61 | 0.8 | 0.79 | 0.66 | 0.69 | 0.57 | 0.58 | 0.74 | 0.82 | 0.64 | 0.68 | 0.57 | 0.61 | 0 | 0.96 | 0.82 | 0.83 |
| S2_RNA-Seq | 0.8 | 0.74 | 0.58 | 0.78 | 0.77 | 0.63 | 0.66 | 0.54 | 0.55 | 0.75 | 0.81 | 0.63 | 0.68 | 0.56 | 0.6 | 0.96 | 0 | 0.81 | 0.82 |
| Unfertilized_Egg_Riboseq | 0.86 | 0.75 | 0.57 | 0.88 | 0.86 | 0.63 | 0.67 | 0.52 | 0.54 | 0.76 | 0.87 | 0.61 | 0.63 | 0.5 | 0.59 | 0.82 | 0.81 | 0 | 0.93 |
| Unfertilized_Egg_RNA-Seq | 0.86 | 0.72 | 0.54 | 0.83 | 0.81 | 0.59 | 0.62 | 0.48 | 0.5 | 0.79 | 0.88 | 0.6 | 0.62 | 0.49 | 0.57 | 0.83 | 0.82 | 0.93 | 0 |

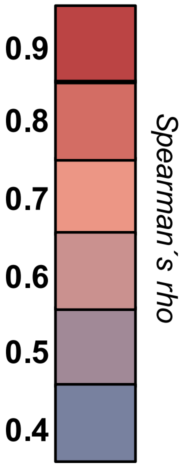

### Supplementary Tables 3

**A** *tal*

| Measure | Stage | Replica | *tal-1A* | *tal-2A* | *tal-3A* | *tal-AA* |
|---|---|---|---|---|---|---|
| | | | *ORF* | | | |
| RNA-Seq RPKM | Early | T | 19.75 | 87.08 | 154.42 | 45.70 |
| | Early | B | 10.28 | 102.47 | 139.96 | 61.70 |
| | Mid | T | 110.55 | 600.49 | 1560.27 | 130.65 |
| | Mid | B | 57.59 | 792.04 | 508.15 | 335.57 |
| | Late | T | 4.06 | 36.12 | 29.80 | 12.15 |
| | Late | B | 1.49 | 24.80 | 40.17 | 9.74 |
| Ribo-Seq RPKM | Early | T | 3.60 | 7.20 | 7.20 | 8.51 |
| | Early | B | 19.53 | 35.30 | 68.72 | 33.46 |
| | Mid | T | 36.94 | 34.89 | 59.52 | 47.02 |
| | Mid | B | 76.78 | 147.40 | 293.56 | 144.52 |
| | Late | T | 4.19 | 4.19 | 12.58 | 9.65 |
| | Late | B | 13.27 | 39.81 | 53.07 | 31.50 |
| TE | Early | T | 5.48 | 12.09 | 21.43 | 5.37 |
| | Early | B | 0.53 | 2.90 | 2.04 | 1.84 |
| | Mid | T | 2.99 | 17.21 | 26.21 | 2.78 |
| | Mid | B | 0.75 | 5.37 | 1.73 | 2.32 |
| | Late | T | 0.97 | 8.62 | 2.37 | 1.26 |
| | Late | B | 0.11 | 0.62 | 0.76 | 0.31 |
| **translation probability** (binomial p) | **Early** | Both | **1.54E-05** | **3.56E-11** | **7.36E-10** | **5.79E-09** |
| | **Mid** | Both | **5.62E-07** | **2.32E-19** | **1.75E-18** | **7.50E-10** |
| | **Late** | Both | 6.57E-01 | 7.44E-02 | **7.06E-06** | 6.45E-01 |
| **translational regulation** (Z-ratio) | **Early-Mid** | Both | 1.22E-01 | 3.26E+00 | 2.93E+00 | 2.05E-01 |
| | **Early-Late** | Both | -5.01E-01 | -3.98E+00 | -4.36E+00 | -1.05E+00 |
| ***Conservation (phyloP)*** | | | 2.5 | 3.1 | 2.8 | 2 |

**B** *scl*

| Measure | Stage | Replica | *scl-A* | *sclB* |
|---|---|---|---|---|
| | | | *ORF* | |
| RNA-Seq RPKM | Early | T | 0.00 | 1.44 |
| | Early | B | 1.24 | 0.75 |
| | Mid | T | 5.94 | 17.24 |
| | Mid | B | 6.63 | 20.20 |
| | Late | T | 6.36 | 3.35 |
| | Late | B | 9.69 | 20.92 |
| Ribo-Seq RPKM | Early | T | 0.00 | 1.08 |
| | Early | B | 0.57 | 0.56 |
| | Mid | T | 5.20 | 50.05 |
| | Mid | B | 2.24 | 11.92 |
| | Late | T | 74.74 | 140.88 |
| | Late | B | 55.62 | 59.71 |
| TE | Early | T | N/A | 0.75 |
| | Early | B | 0.46 | 0.74 |
| | Mid | T | 0.87 | 2.90 |
| | Mid | B | 0.34 | 0.59 |
| | Late | T | 11.75 | 42.01 |
| | Late | B | 5.74 | 2.85 |
| **translation probability** (binomial p) | **Early** | Both | 3.00E-01 | 1.00E+00 |
| | **Mid** | Both | **4.00E-03** | 3.40E-02 |
| | **Late** | Both | **7.20E-52** | **3.84E-08** |
| **translational regulation** (Z-ratio) | **Early-Mid** | Both | 1.70E-01 | 2.70E-01 |
| | **Early-Late** | Both | 2.03E+00 | 3.30E+00 |
| ***Conservation (phyloP)*** | | | 2.6 | 2.9 |

### Supplementary Table 3

| ORF class | | Transcribed ORFs | | Total |
|---|---|---|---|---|
| **Canonical** | 14469 | 16273 | 16332 | 17280 |
| **shortCDS** | 402 | 436 | 471 | 490 |
| **uORF** | 8454 | 9858 | 9069 | 11710 |
| | | **Ribo-Bound** ORFs | | |
| **Canonical** | 12732 | 15650 | 15544 | 16858 |
| **shortCDS** | 295 | 386 | 405 | 443 |
| **uORF** | 5691 | 6669 | 5416 | 8346 |
| | | **Translated** ORFs | | |
| **Canonical** | 12159 | 14300 | 14459 | 15956 |
| **shortCDS** | 254 | 270 | 309 | 355 |
| **uORF** | 1046 | 653 | 617 | 1462 |
| | *Early* | *Mid* | *Late* | *Embryo* |
